# Iguanas from above: Citizen scientists provide reliable counts of endangered Galápagos marine iguanas from drone imagery

**DOI:** 10.1101/2024.02.09.579637

**Authors:** Andrea Varela-Jaramillo, Christian Winkelmann, Gonzalo Rivas-Torres, Juan M. Guayasamin, Sebastian Steinfartz, Amy MacLeod

**Affiliations:** Institute of Biology, Molecular Evolution and Systematics of Animals, University of Leipzig, Leipzig, Saxony, Germany; 3Diversity, Quito, Pichincha, Ecuador; Eberswalde University for Sustainable Development, Eberswalde, Brandenburg, Germany; Laboratorio de Biología Evolutiva, Instituto Biósfera, Colegio de Ciencias Biológicas y Ambientales COCIBA, Universidad San Francisco de Quito USFQ, Calle Diego de Robles s/n y Pampite, Cumbayá, Quito, Pichincha, Ecuador; Galápagos Science Center, GSC, San Cristóbal, Galápagos, Ecuador; Wildlife Ecology and Conservation, Gainesville, University of Florida, FL, United States of America

**Author notes:** **Correspondence** Amy MacLeod.

**Keywords:** Citizen Science, drone imagery, wildlife monitoring, Zooniverse, wildlife surveys

## Abstract

Population surveys are vital for wildlife management, yet traditional methods often demand excessive time and resources, leading to data gaps for many species. Modern technologies such as drones can facilitate field surveys but may also increase data analysis challenges. Citizen Science (CS) can address this issue by engaging non-specialists for data collection and analysis. We evaluated CS for population monitoring using the endangered Galápagos marine iguana as a case study, assessing online volunteers’ ability to detect and count animals in aerial images. Comparing against a Gold Standard dataset of expert counts in 4345 images, we explored optimal aggregation methods from CS inputs, considering image quality and filtering data from infrequent and anonymous participants. During three phases of our project — hosted on the Zooniverse platform — over 13,000 volunteers made 1,375,201 classifications from 57,838 aerial images; each being independently classified 20 (phases 1 & 2) or 30 (phase 3) times. Volunteers achieved 68% to 94% accuracy in detecting iguanas, with more false negatives than false positives. Image quality strongly influenced accuracy; by excluding data from suboptimal pilot-phase images, volunteers counted with 90% to 92% of accuracy. For detecting presence or absence of iguanas, the commonly used ‘majority vote’ aggregation approach (where the answer selected is that given by the majority of individual inputs) produced less accurate results than when a minimum threshold of five (from the 20/30 independent classifications) was used. For counting iguanas, HDBSCAN clustering yielded the best results. Excluding inputs from anonymous and inexperienced volunteers decreased accuracy. We conclude that online volunteers can accurately identify and count marine iguanas from drone images, though a tendency to underestimate warrants further consideration. CS-based data analysis is faster than manual counting but still resource-intensive, underscoring the need to develop a Machine Learning approach.

## Introduction

Population surveying and species monitoring are central tasks in conservation work, since the information gathered forms the foundation of wildlife management plans. However, for many species, collecting such data by traditional means often requires more resources and time than are available. At present, of the 81% of vertebrate species assessed by the IUCN, around 14% are data deficient (DD) (IUCN, 2024), with unknown population status and distribution range being the key reasons for a DD listing in many groups, which includes amphibians, reptiles, and mammals (Bland et al., 2017). Modern approaches employing new technologies — such as drones for aerial surveys — are greatly reducing monitoring effort and surveyor risk whilst simultaneously allowing access to remote regions (Ezat et al., 2018; Lee et al., 2019; Monks et al., 2022; Ratcliffe et al., 2015). However, with the rise in popularity of image-based approaches, comes an increased burden for data analysis; this presents a significant challenge for conservation practitioners due to the time and resource constraints inherent in such work. This bottleneck can be eased through the development of novel analysis approaches in order to ensure the long-term future of applying image-based methods.

One increasingly popular approach involves mobilizing non-specialist help in the form of Citizen Scientists (CS; also referred to as volunteers). Whilst there is already a myriad of large-scale successful projects — such as iNaturalist (www.inaturalist.org), Wildbook (Berger-Wolf et al., 2017), Coral Watch (Marshall et al., 2012), and e-Bird (Sullivan et al., 2014) where CS collect data — engaging volunteers for data analysis is relatively less developed. In the two-plus decades that such projects have been running, online citizen science projects have already demonstrated that volunteers can reliably identify target species and count individuals from photographs with minimal training (Cox et al., 2015). By such ‘crowd-sourcing’ data analysis (popularly known as ‘the wisdom of crowds’), researchers can massively reduce the amount of specialist time and effort needed to obtain results (e.g. Swanson et al., 2015). To this end, we herein explore the application of CS-powered data analysis for surveys of the endangered marine iguana (*Amblyrhynchus cristatus;* MacLeod et al., 2020) — a charismatic and well-known endemic species of the Galápagos.

For nine of the 11 subspecies of marine iguana (Miralles et al., 2017), population-size data are severely lacking, due in large part to the immense logistical challenges inherent in surveys of this species (MacLeod & Steinfartz, 2016). Given the unprecedented levels of anthropogenic change (e.g., introduced species, climate change, marine pollution) on the Galápagos (Alava et al., 2023), the need for better information is urgent. In order to address this data gap, we employed a recently validated boat-based surveying method that uses drones to collect aerial images of the marine iguana colonies (Varela-Jaramillo et al., 2023) from its entire known distribution range. However, this approach generates a large image-based dataset that would typically require immense specialist effort to analyse, delaying the acquisition of results and reducing the future use of this approach. In seeking a solution, in 2020 we created a project on Zooniverse.org; the largest platform for online citizen science projects (Simpson et al., 2014). In the project — named “Iguanas from Above” — we placed a set of photographs (smaller slices of orthomosaic images created from the raw drone images) from a selection of locations. Each image was shown to a predetermined number of independent volunteers, who were asked to mark iguanas within the photograph; this task is known as a “classification”. Once all images were classified, the resulting raw dataset would then be “aggregated”; a procedure that obtains a consensus of results for each image from the individual inputs of each volunteer. Crowdsourcing data analysis has much to offer for conservation projects since it can greatly reduce the expert effort required for analysis whilst simultaneously engaging the public in conservation work. However, in comparison to extensive work done in projects where volunteers perform data collection, analysis and validation of CS-sourced wildlife counts are currently lacking. Having now run three successive phases of our Zooniverse project (each covering data from one annual field season), we herein use this as a case study for the use of CS in the analysis of image-based datasets for the purpose of population-size estimation in wild populations in the Galápagos.

Within this work, we aim to address the following questions:

1. **In what ways can the results from the raw CS dataset be filtered to improve agreement between citizen scientists and experts?** Within this query, we addressed the following sub questions: (a) does removing inputs from anonymous or infrequent volunteers improve the outcome? And (b): does increasing the number of times an image is independently classified (‘classifications required per image’) increase count accuracy?
2. **Which aggregation method used produce results closest to those of experts?** As we have up to 30 separately entered counts per image (one from each volunteer classification), it is necessary to summarise the outcome on aggregate. Here we test traditional measures such as mean, median and mode, as well as some approaches that integrate the coordinates of iguanas — as marked by the volunteers — as an additional consideration.
3. **Which aspects of the images presented to citizen scientists are important when considering accuracy of results?** For this, we investigated the phase analysed; the quality of the image in terms of sharpness, blur, lighting, etc.; and the number of individuals present in the image as potential factors.
4. **Can citizen scientists accurately identify and count marine iguanas on a given aerial image?** This is — in essence — our overarching question. To answer it, we compare aggregated volunteer counts to those of ‘experts’ — I.e. professional scientists trained for marine iguana detection in aerial images. In accordance with previous studies across diverse systems and taxa (e.g. Cosentino & Gibbs, 2022; Kosmala et al., 2016; Lintott et al., 2008; Swanson et al., 2016), we define CS data as accurate when there is an agreement of > 95% with experts for object detection (i.e. correctly identifying the presence or absence of marine iguanas in the image) and > 80% for object counts (i.e. matching the expert count for the number of marine iguanas in the image). If sufficient agreement is found, the overall goal is to utilize CS inputs for the estimation of population size at key colonies of marine iguanas.

## Methods

### Building the citizen science project

We collected the aerial images used for this project using commercial drones (DJI Mavic 2 Pro), flown from land and boats along the rocky coastline of several marine iguana colonies in the Galápagos Archipelago during three successive field seasons from 2020 to 2022 (Fig. 1). From these, we created orthomosaics (2D-georeferenced maps) using the software Agisoft Metashape; for full details on image collection and analysis protocols, see Varela-Jaramillo et al. (2023). We ‘sliced’ the orthomosaics using Adobe Photoshop to create individual images of 1000x1000 pixels on average (resulting image size: up to 1 MB). We did this in an attempt to standardize the size of the individual iguanas in the images to aid recognition; standardizing was necessary because the size of iguanas in the image depends on both the height at which we flew the drones (20-30 m altitude; which due to complex topography could not be accurately measured) and the subspecies of the marine iguana (who vary significantly in body size on different islands).

**Figure 1.**
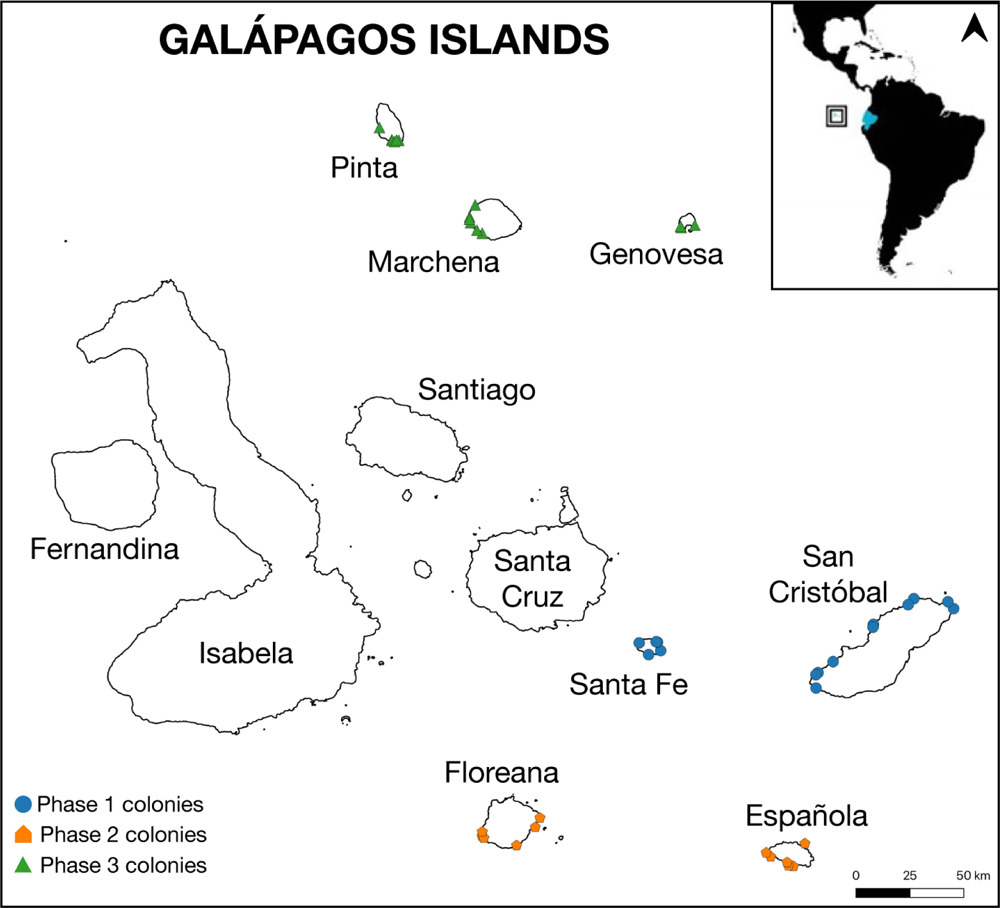
Map of the Galápagos Islands. The coloured points represent all the colonies surveyed for this study. The three phases analyzed covered seven of the major islands. The square in the upper right shows the geographic location of the Archipelago, ∼1000 km from the coast of continental Ecuador.

We created our project on the Zooniverse platform in English with additional instructions in German, and phases one and two were also available in Spanish and French. Our workflow for the project requires volunteers to complete three tasks for each image classification (Fig. 2): 1) identify presence or absence of marine iguanas in the image; 2) mark individuals of marine iguanas, distinguishing adult males and reproductive groups (leks) from ‘others’ (females, sub-adult males, and juveniles) when possible. The category ‘partial iguana’ was an addition made to avoid double-counting of occasional individuals that were bisected at the edge of the image during slicing; and 3) identify and count individuals of cohabiting species which included sea lions (*Zalophus wollebaeki*), crabs (*Grapsus grapsus*), Green Turtles (*Chelonia mydas*) sea birds, plants, and algae (various species), as well as plastic objects (bottles and fishing gear). We provide a tutorial and a ‘Field Guide’ for species identification and a message board for discussion; this enables volunteers to interact with researchers and each other regarding image classification, and also provides a space for ongoing discussion on Galápagos wildlife and conservation matters.

**Figure 2.**
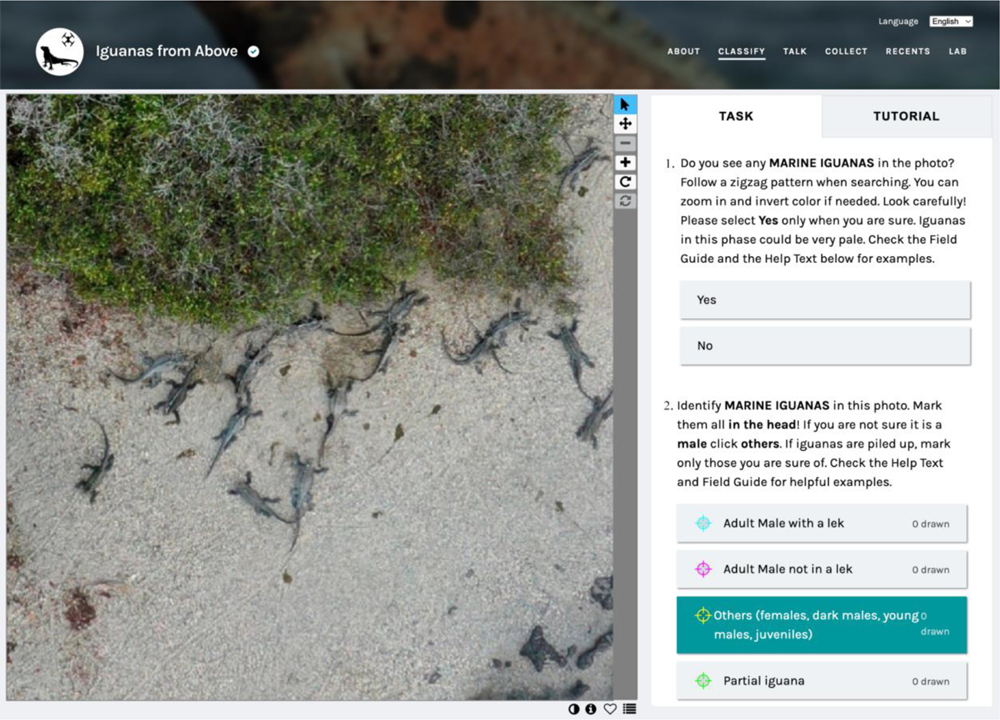
Iguanas from Above citizen science web portal in Zooniverse.org. An example tile with two tasks presented to citizen scientists: Task 1, regarding identification of marine iguana presence or absence, and Task 2 where volunteers mark the number of individuals present in the image.

To date we have launched four phases of the Zooniverse project, each following an annual fieldtrip undertaken between 2020 and 2023, during which we surveyed selected islands within the mating season of the marine iguana. For this study, we analysed the first three complete phases. Phase 1 began in August 2020, and comprised 24,373 images from San Cristobal and Santa Fe islands surveyed as a pilot project in January 2020. The three subspecies covered have medium to large body sizes (Miralles et al., 2017) and relatively low densities. Phase 2 — launched in February 2022 — included 9,097 images from Española and Floreana islands, collected in January 2021. This phase featured the more colourful and abundant “Christmas iguana”, which has a medium to large body size and is renowned for the turquoise coloration displayed by males in the mating season. Phase 3 was launched in July 2022 with 24,368 images from the northern islands of Genovesa, Marchena, and Pinta, surveyed in December 2021. Marine iguanas on these islands are rare, small-bodied, highly cryptic against the rocky substrate, and males appear to lack the coloration seen on other islands in the pairing season. Two sites from Phase 2 were also resurveyed in 2021 and re-launched here.

We circulated each image among a predefined number of independent volunteers, whose input on an image is referred to as a classification. We required 20 classifications per image for phases 1 and 2, and 30 classifications for phase 3. The extra 10 classifications on the latter phase was intended to test whether more classifications per image would improve the accuracy of CS outputs.

To increase volunteer participation, we undertook promotional activities including press releases, newsletters, inviting inputs from schools, universities and companies via webinars, social media and blog posts; as well as participating on the citizen science month event promoted by SciStarter (https://scistarter.org/iguanas-from-above) and being a featured Zooniverse project. We also ran a competition to award the best classifiers in different categories as a means to motivate the volunteers.

### Building the Gold-Standard dataset for CS-data analyses

We used RStudio version 2023.09.1+494 (RStudio Team, 2023) and python version 3.10.13 (Van Rossum & Drake, 2009) with the library Pandas version 2.1.2 (The Pandas Development Team, 2020) for data frame management, ensuing analyses and plotting of results. We downloaded CS-classifications from the Zooniverse platform and used the Panoptes Aggregation python package (https://aggregation-caesar.zooniverse.org/Scripts.html) to extract and summarise (‘aggregate’) data from Task 1 (marine iguana presence/absence – question-type data) and Task 2 (marine iguana counts – point-type data). We randomly selected around 5-10% of the images per phase as a Gold-Standard (GS) dataset; these images were chosen to cover a range of challenge including a variety of iguana body sizes and coloration, both high and low density colonies, and from various fieldwork years (since quality of images improved over the course of the project). In total, this included 4345 images from 30 colonies selected from 7 islands; 2733 images from phase 1, 456 images from phase 2 and 1156 images from phase 3. Three people from the research team (henceforth referred to as ‘experts’) analysed the GS datasets and generated consensus results (henceforth simply called ‘expert’ data) for presence/absence; number of iguanas in the image, and a judgement on quality of the image (simply “good” or “bad”; Fig 3a). Image quality judgement was based on consideration of: sharpness of camera focus, light levels within the image, image blur due to camera movement, image ‘smear’ due to mosaicking artefacts, and complexity of the substrate on which the animals occur. The expert consensus count was taken as the highest number of iguanas agreed between at least two of the three counters. We then compared this ‘expert data’ to the CS data for each phase, and for all images together. The GS dataset has been made available in Dryad (https://doi.org/10.6084/m9.figshare.25196306).

**Figure 3.**
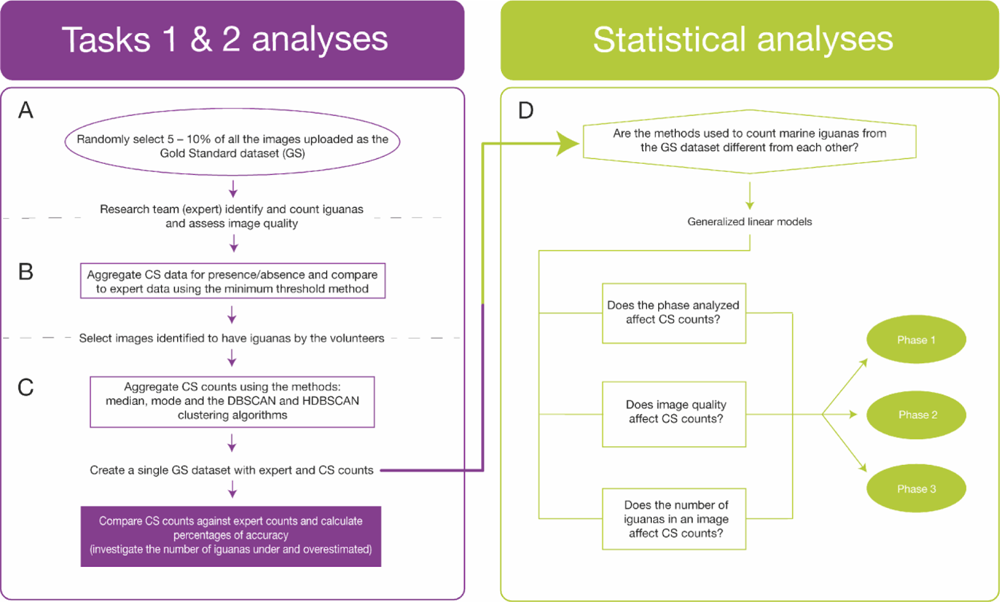
Workflow used to analyse citizen science data obtained for the project. Analyses to test CS accuracy in Tasks 1 and 2 and the statistical analysis to investigate potential factors affecting their accuracy.

### Marine iguana presence or absence analyses

In the online task, each volunteer is asked whether marine iguanas are present (‘yes’) or absent (‘no’) in the image; therefore, for each image in our dataset, we obtained 20 (phase 1 & 2) or 30 (phase 3) answers. A commonly used approach for aggregating (i.e. summarising) results where multiple volunteers give input is the ‘simple plurality algorithm’ (also known as ‘majority vote’; Swanson et al., 2015). This is where the answer selected by 50% plus one of the volunteers for any given task is accepted as the result for this image. For example, in our case, where 20 volunteers classified an image and 11 or more selected ‘yes’ to indicate presence of an iguana, the result would be ‘iguana(s) present’. We sought to find >95% agreement with the expert classifications, which we define as the minimum agreement level deemed accurate for our purposes. We were interested in testing the performance of the ‘majority vote’ approach, and specifically sought to test whether a smaller number of volunteers could also give accurate results. For this purpose, we calculated the agreement between CS and expert 11 times for each image; giving an agreement level for the case where one of 20 volunteers selected ‘yes’, then two of 20 and so on up to 11 of 20. The aim was to find a ‘minimum threshold’ for the smallest number of volunteers indicating iguana presence in (95%+) agreement with the experts (Fig. 3b).

We also analysed if removing anonymous (i.e. users not logged into the platform) inputs increased volunteer accuracy in identifying marine iguanas, to explore whether project loyalty and volunteer (in)experience might have an effect on the reliability of inputs. Likewise, we tested how removing inputs from volunteers with 10 or fewer classifications affected accuracy, since we considered that these infrequent participants have less experience at the task and thus may be less skilled.

### Marine iguana counting analyses

We analysed whether, after selecting yes for marine iguana presence, volunteers were able to detect all the marine iguanas present on the image. In this task, volunteers added marks to the image to indicate where they detected the iguanas (Fig. 2). To analyse these outputs, we first selected all images where iguanas were present (as obtained using the ‘minimum threshold’ rule) and aggregated the volunteer counts for each (Fig. 3c). We calculated two statistical metrics to summarise these counts: the median and the mode. The median searches for the value in the middle of an ordered data sample, while the mode seeks for the most repeated value within the sample. These metrics are appropriate because they eliminate the outlier effect we anticipated to find from occasional volunteers not fully engaged in their task. Additionally, we tested the Density-Based Spatial Clustering of Applications with Noise algorithm (DBSCAN) and the Hierarchical Density-Based Spatial Clustering of Applications with Noise algorithm (HDBSCAN), to collate the volunteer annotation marks into spatial clusters of points within the image. The amount of clusters then represents the amount of iguanas present in the image (Pedregosa, et al., 2011) (See Supplementary Fig. S1 and Supplementary Methods for further details).

From these metrics, we investigated volunteer accuracy by comparing counts between experts and CS for each image. Specifically, we looked for: the number of images where volunteer counts were in 100% agreement with expert counts; the number of images where volunteers counted fewer iguanas than the experts (underestimation); and where CS counts were higher than experts (overestimation). Finally, we summed all GS aggregated counts by method used, and compared these to the expert counts, calculating the percentage of agreement (‘accuracy’), as well as the number of iguanas missed and over counted.

### Statistical analyses regarding volunteer accuracy when counting marine iguanas

We statistically explored — using generalized linear models with quasi-poisson corrections — how counts obtained in GS images related among the different methods used to obtain them (expert consensus, median, mode, and HDBSCAN; DBSCAN clustering analysis was excluded from statistical analysis due to poor initial results). We ran ANOVAs, and calculated the Estimated Marginal Means (R package emmeans; Lenth et al., 2023) to test the significance of pairwise comparisons among all methods (Fig 3d). Further, a logistical regression was performed to find which CS data aggregation method best fit the expert data.

We also investigated the effect of three factors on volunteer counts using the same statistical methods, these were: phase analysed; image quality (see discussion on GS within methods for criteria); and the number of iguanas present on the image (for this we defined three categories based on the data distribution: Low: 1-5, medium: 6-10, and high: >10 marine iguanas; see Supplementary Fig. S2).

### Citizen scientists’ experiences

In order to better understand the volunteers’ experiences, we undertook a short online survey. This included questions referring to the number of images classified, factors affecting their motivation to participate in the project, and differences they perceived between the phases (see Supplementary Methods for details). These results were explored to better understand the relationship between volunteer perceptions and our results, and were also used for the purposes of improving our ongoing CS work.

## Results

### Volunteer participation insights and filtering

The three phases included in this study were completed in September 2023. At this point, over 10,000 registered volunteers and 3500 unregistered (anonymous) volunteers (13,988 in total), had made 1,375,201 classifications from 57,838 aerial images. The majority of volunteers (86%, n = 12,099) made less than 50 classifications, yet some classified thousands of images (174 volunteers each classified over 1,000 images). These results agree with those provided in the survey carried out by the volunteers (see Supplementary Fig. S3), where most respondents reported to have entered 100 or fewer classifications. A small number of ‘super volunteers’ made a large input: our top 10 contributors represented 18% (n = 244,654) of the total classifications.

Logging-in to classify images is a Zooniverse recommendation, but it is not mandatory, and we found overall that 15% (n = 207,942) of our classifications were performed by participants who were not logged-in (3753 volunteers). Even when a user is not logged in, Zooniverse gives them an identification, which is registered as “not-logged-in+IP number”. An important amount of data (16%) in the GS dataset was generated by anonymous volunteers; when this data was excluded, the accuracy within the ‘iguanas present’ images decreased by 9% in phase 1, 2% in phase 2, with no reduction in phase 3. Therefore, all classifications — including anonymous inputs — were retained for the analysis.

### Aggregation methods: minimum threshold for accurate CS counts

Considering all images within the GS dataset (i.e. those with and without iguanas) the highest level of accuracy was found when 5 or more volunteers (from the 20 or 30 that classified each image) indicated the presence of an iguana. This finding was consistent across all phases of the project (Fig. 4a). Agreement levels across each phase were: phase 1 (98%; n = 2733); phase 2 (97%; n = 456); and phase 3 (97%; n = 1156). Accuracy decreased when the majority vote method was used (phase 1 = 97%; phase 2 = 93%; and phase 3 = 95%; see supplementary Fig. S4); therefore, our results suggest that requiring more repeated classifications of an image does not increase CS accuracy. Since the vast majority (90%) of our images were blank (i.e. did not contain an iguana), further analysis was undertaken to assess the effect of this skew on the overall dataset. For this, we applied the minimum threshold of five volunteers as the aggregation method.

**Figure 4.**
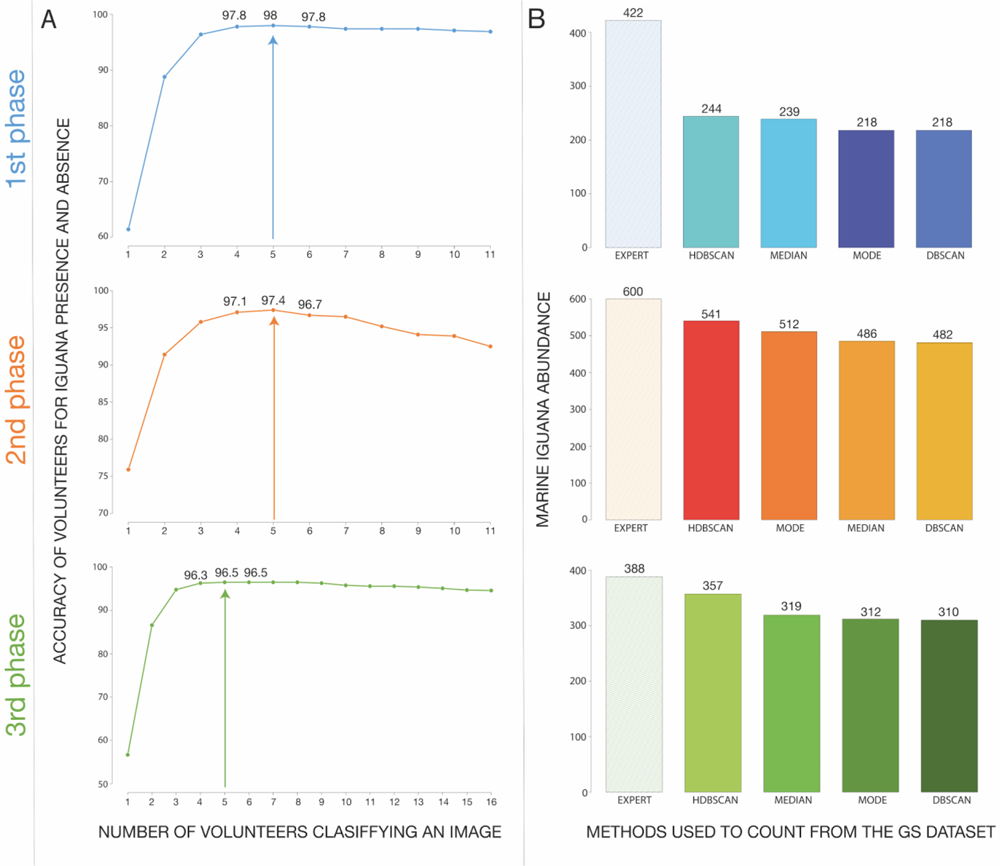
Results from Task 1 and 2. A) Minimum volunteer threshold results for all phases analysed, and B) total marine iguana counts obtained by the experts, across all methods used to aggregate CS data from multiple inputs.

When experts did not detect any iguanas in an image, we classified the image as ‘absent’. Using only these ‘absent’ images, we compared the expert results to the CS data. Here we classified the CS outcome as correct if fewer than five volunteers indicated an iguana in an image recorded as ‘absent’ by experts. In this, we report an average of 99.6% of agreement between experts and CS across the three phases. In images where iguanas were identified by experts (10% of images overall), we classified these as ‘present’. For this, we deemed the CS outcome as correct if five or more volunteers indicated an iguana in an image recorded as ‘present’ by experts. Volunteers classified these images correctly in 68% of images in phase 1 (n = 150), 94% in phase 2 (n = 179) and 70% in phase 3 (n = 116), meaning that false negatives varied by phase, and were more likely than false positives.

### Aggregation methods: Marine iguanas as counted by the volunteers

Of the 4345 images in our GS dataset, volunteers found iguanas to be present in 445 (10%) of the images when applying the minimum threshold rule as outlined above, thus allowing comparison to expert counts within this subset of images. For each image, we obtained the aggregated count of the 20/30 volunteers who assessed the image using various metrics for this comparison. When combining results from all phases, we found that volunteer counts matched those of the experts in 94.2% of the images when using the median, in 94.5% with the mode, 94.6% with the HDBSCAN clustering analysis and 92.1% with the DBSCAN approach; though this varied by phase from 80 to 97% (see Supplementary Table S1). We found significant differences among the outcomes of the various methods used to count marine iguanas (comparing expert counts to CS data aggregated using the median, mode, and HDBSCAN; anova test; df = 3, *P* < 0.01), with HDBSCAN obtaining the best results from the CS data.

Considering only images where iguanas were present, we found that volunteer counts were significantly lower than those of experts, when the aggregated counts used both the median (emmeans test; *z*. ratio = 3.33, *p* = 0.0047) and the mode (emmeans test; *z*. ratio = 3.38, *p* = 0.0040). However, expert counts were similar to those obtained using the HDBSCAN approach (emmeans test; *z*. ratio = 2.40, *p* = 0.767). We did not find significant differences between CS results calculated between the median and the mode (emmeans test; z. ratio = 0.047, p = 1.0). Logistical regression analysis obtained the highest fit value with the expert counts for the HDBSCAN approach (Nagelkerke R-quadrat: 0.912), followed by the median (Nagelkerke R-quadrat: 0.885), and the mode (Nagelkerke R-quadrat: 0.856 (see Supplementary Table S2).

Finally, we compared the sums of all counts across the whole project and by individual phase. Total expert counts per phase were: 422 iguanas in phase 1,600 in phase 2, and 388 in phase 3. For total SC counts, HDBSCAN obtained the highest agreements in all phases: 58% in phase 1, 90% in phase 2 and 92% in phase 3. These results were followed by the median and the mode. Although phase 2 showed lower accuracies in terms of number of images where CS and expert were the same (see Supplementary Table S1), underestimation of iguanas was higher in phase 1 (Table 1; Fig. 4b).

**Table 1.**
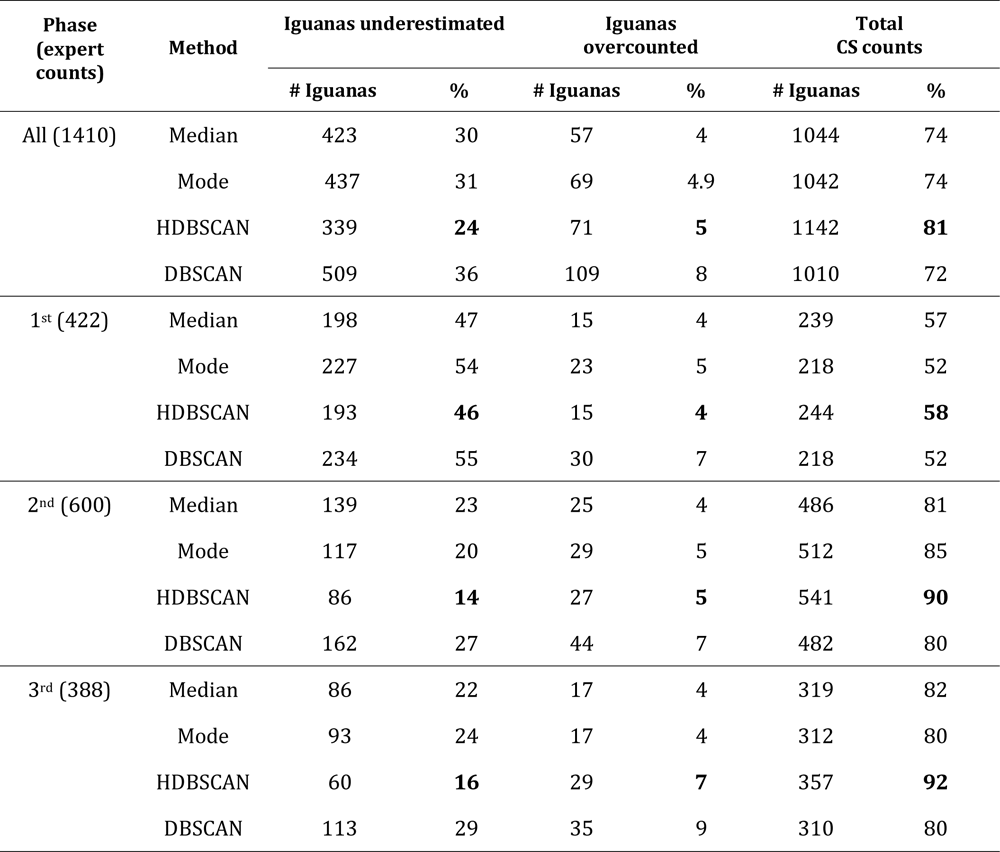
Number of images where volunteer counts (‘CS’) were the same as experts; those where volunteers counted less than the experts (underestimation) and those where volunteers counted more. Bold values represent best results of the aggregation methods.

### Factors affecting volunteer classifications

We also investigated whether the factors: phase of project; quality of the image (classified as “good” or “bad” by the experts); and number of iguanas present in the image (in three categories of low, medium or high; see methods for further details) potentially affected the volunteers-counts. When we added the factor “phase” to the model, we found significant differences among the methods (anova test; df = 3, *P* < 0.001). Specifically, expert counts were significantly higher than CS-counts as calculated with the median (emmeans test; z. ratio = 3.44, *p* = 0.0284) and the mode (emmeans test; z. ratio = 3.49, *p* = 0.0242), but not when HDBSCAN was used (emmeans test; z. ratio = 2.47, *p* = 0.3532). However, when we investigated CS count response as analysed by phase independently, we found different results. In phase 1, the expert counts were significantly higher than all the methods used to obtain CS-counts: median (emmeans test; z. ratio = 3.905, *p* = 0.0005), mode (emmeans test; z. ratio = 4.518, *p* < 0.0001) and HDBSCAN (emmeans test; z. ratio = 3.725, *p* = 0.0011). Interestingly, in phase 2 we found non-significant differences among the expert and all the aggregating methods (anova test; df = 3, p = 0.5175). Likewise, in phase 3 (anova test; df = 3, p = 0.5059). Apparently, phase 1 greatly influenced not only the results of the model, but also the volunteer counts (Fig. 5)

**Figure 5.**
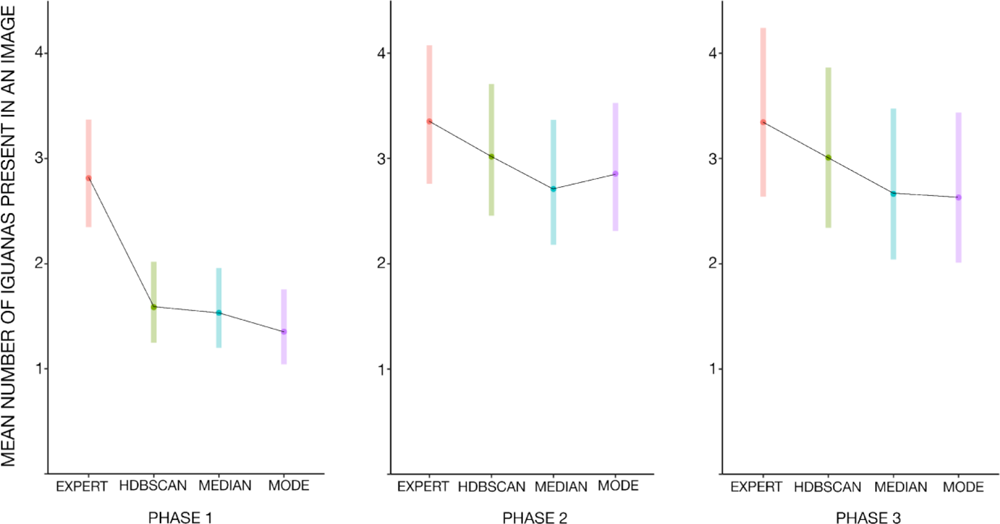
Plot of the generalized linear model results when all the methods used to count marine iguanas from the GS images were compared and analysed by phase independently. The black lines join the means of all methods, and the bars represent the data variation of each method (intervals of confidence).

When we added the factor “image quality” to the model, we found significant differences among the methods (anova test; df = 3, *P* = 0.0013). Expert counts were significantly higher than the CS counts, both in images with good and bad quality, when using the median (emmeans test; z. ratio = 3.333, p = 0.0194) and the mode (emmeans test; z. ratio = 3.379, p = 0.0167), but not different when using the HDBSCAN method (emmeans test; z. ratio = 2.399, p = 0.2418). In general, there were no significant differences between the average number of iguanas per image in relation to image quality (anova test; df = 1, p = 0.3635). However, when we investigated the influence of image quality on CS-counts as analysed by each phase independently, we found conflicting results. In phase 1, the expert counts were significantly higher than all the methods on images of both good and bad quality: median (emmeans test; z. ratio = 3.920, p = 0.0023), mode (emmeans test; z. ratio = 4.536, p = 0.0002) and HDBSCAN (emmeans test; z. ratio = 3.741, p = 0.0045). Here, the differences among the average number of iguanas per image significantly differed among quality (anova test; df = 1, P = 0.0332), where images containing more iguanas were typically of poor quality. On the other hand, when analyzed separately, differences in phase 2 are no longer significant among metrics (anova test; df = 3, p = 0.5173), or between image quality categories (anova test; df = 1, p = 0.8568). This was also true for phase 3 across metrics (anova test; df 3, *p* = 0.5073) and image quality (anova test; df =1, *p* = 0.5335) (see Supplementary Fig S5).

Finally, we found that the number of marine iguanas present on the image did not greatly influence the results. Expert counts were significantly higher than the CS counts with all methods used in all categories of abundance (see Methods): median (emmeans test; z. ratio = 4.674, *p* = 0.002), mode (emmeans test; z. ratio = 4.702, *p* = 0.002) and HDBSCAN (emmeans test; z. ratio = 3.363, *p* = 0.0371). However, once again, this result differed when the analysis was performed independently for each phase. In phase 1, significant differences between the expert and CS data from various metrics remain regardless of the number of iguanas present: median (emmeans test; z. ratio = 4.577, p = 0.003), mode (emmeans test; z. ratio = 5.161, *p* <0.001) and HDBSCAN. Conversely — as found with the image quality analysis — there were no significant differences between the expert and CS counts in all categories of abundance for both phase 2 (anova test; df = 3, *P* = 0.087) and phase 3 (anova test; df = 3, *P* = 0.2527) (see Supplementary Fig S6). In conclusion, volunteers can also count accurately even when the number of iguanas on an image is high, but where image quality is concurrently poor — a scenario more frequently encountered in phase 1 — CS accuracy decreases.

### Results of Volunteer Survey

We received 110 inputs in our volunteer survey. In terms of perceived difficulty in classifying images, the key issues reported were image blur, poor camera focus, and similarity between substrate (typically dark rocks) and the iguanas (see Supplementary Fig. S7a). Few noted that (too) many iguanas was a problematic feature of the images, (see Supplementary Fig. S7b). In terms of engagement, a lack of iguanas in the images (i.e. ‘blank’ images) was the most important factor in decreasing motivation, followed by poor image quality, similarity of substrate and object, and repetitiveness of the task (see Supplementary Fig. S8).

## Discussion

### Which aggregation method used produces results closest to those of experts?

Remarkably, we found that the simple plurality algorithm (‘majority vote’; Cardoso et al., 2020; Swanson et al., 2015) — frequently used to aggregate data from multiple volunteers — did not produce the most reliable data. Instead, in the task of detecting presence or absence of iguanas within an image, we found that the answer selected by five volunteers or more (from 20 or 30 total classifications) — which we refer to as the minimum threshold — was the most likely to be correct. This was somewhat unexpected, since previous work has shown that accuracy increased asymptotically with number of classifications per image (Swanson et al., 2016); however, other projects hosted by Zooniverse have found outcomes more similar to ours (Egna et al., 2020; Hennon et al., 2015; Lawson et al., 2022). We suspect that for some challenging images, highly skilled volunteers were required to identify the iguanas, and that these volunteers were relatively rare. Therefore, if we applied the majority vote rule, we would require 11 of 20 volunteers who viewed each image to identify the iguana, whereas using a minimum threshold of five would require just five of 20, giving relatively more weight to the highly skilled ones. It may be that each project — with its distinctive dataset and related set of challenges — may require exploration to find the best minimum threshold of classifications for aggregation, at least in cases where the majority rule does not produce the required level of accuracy.

In images where experts found no iguanas (‘absent’ or ‘blanks’), CS results were in 99.6% agreement; thus our results show a very low instance of false positives (i.e. volunteers indicating the presence of an iguana in an image when experts did not detect one). This tendency can vary among projects; for instance, in Snapshot Serengeti, volunteers rarely produced false positives (Hines et al., 2015), but in the “Año Nuevo Island Animal Count”, volunteers were more likely to detect non-existent individuals than they were to miss animals (Wood et al., 2021). Although the overall 97% accuracy for presence/absence task is superficially impressive, further analysis is warranted. This is because the majority of our images were blank — i.e. did not contain an iguana, which skews the data. The majority of the 3% error stems from analysis of images where iguanas were present, which was only 10% of the overall images. Within this ‘presence’ data subset, the error rate — so called ‘false negatives’ — is much higher (from 6 –32%); meaning that in our case, volunteers were more likely to miss marine iguanas on the images. This would lead to an underestimation of iguanas by CS.

In terms of counting iguanas, we tested various metrics for summarising (‘aggregating’) multiple volunteer inputs against the expert counts to see which produced most similar results. In this, we found no significant differences among the methods used, although the HDBSCAN obtained the highest accuracies overall in counts for all phases, and obtained the statistical best fit to the expert counts. In contrast to traditional metrics like median and mode, this approach takes into account the digital marks made by volunteers on the images when locating the iguanas. These marks are grouped into clusters, with each representing an individual animal if identified by a minimum number of volunteers — in this case five, using the minimum threshold rule. It seems that use of this additional spatial information may explain the superior performance of the result. Moreover, this positional information of individuals could prove useful for addressing other questions, such as those related to behaviour or habitat use (Wood et al., 2021), as well as providing training data for Machine Learning approaches (Jones et al., 2020). We tested both HDBSCAN and the related approach DBSCAN, and found that the former was not only easier to use — because it required fewer parameters to be set — but also obtained better clustering outcomes (see supplementary information for a deeper discussion on this point). Employing the HDBSCAN clustering analysis, we found that total CS-counts for phase 2 and 3 were 90% and 92% accurate when compared to experts. When the CS counts per image were analysed statistically, we found they were not significantly different to those of the experts. However, the difference between results from HDBSCAN and the median was relatively minor, and this simpler metric may be sufficient for many purposes.

### In what ways can the results from the raw CS dataset be filtered to improve agreement between citizen scientists and experts?

In addition to testing various aggregation approaches we also investigated the impact of filtering input from anonymous and ‘inexperienced’ volunteers (i.e. those with 50 or fewer classifications) from the dataset. We found that anonymous (i.e. users not logged into the platform) contributed around 15% of our total classifications, while inexperienced volunteers — those with fewer than 50 classifications — represented (76%) of all the participants and made 132004 classifications (9%). Eliminating input from such participants has been helpful in some citizen science projects (Dickinson et al., 2010), however — as noted by Swanson et al. (2016) —when having multiple independent classifications per image (crowdsourcing), eliminating classifications often means discarding significant levels of volunteer effort, as well as potentially valuable information. In our case, removing anonymous inputs reduced the overall number of classifications, which in turn resulted in a reduction in accuracy. Worth noting, was that some anonymous volunteers undertook several thousands of classifications (up to 11,606), indicating that being logged in — and thus potentially receiving recognition for inputs — is not necessarily an indication of engagement with a project.

### Which aspects of the images presented to citizen scientists are important when considering their accuracy?

Particularly in phase 1, we found that poor image quality significantly affected the results — this was most obvious on images collected from El Miedo in Santa Fe island. El Miedo was also coincidentally the colony with the highest density within that phase, and is thus the primary reason for the higher values of iguana underestimation here. This colony was part of our pilot phase, where we had limited experience with image collection, and thus the drone protocols were not optimal in terms of altitude and image-overlap. The resulting images were of comparably lower quality and the iguanas are relatively small within the images. Moreover, in this initial phase, we added a watermark to each image, which negatively influenced the visibility of objects within the image. Our image collection procedure has since significantly improved; the impact of this was evidenced by the results obtained in phases 2 and 3. For this reason, we argue that the two later phases are more representative of the approach, and thus should be used here to validate the method.

In addition to image quality, the general predominance of ‘blank’ images within the dataset was also an issue. This scarcity of images in which iguanas were present undoubtedly reduced opportunities for volunteers to ‘learn by doing’. This issue is common among projects where focal objects are rare, too small, or too similar to the background (McShea et al., 2016; Wood et al., 2021) and may have contributed to the false negative error rate, particularly in phase 1 where the density of iguanas on one of the covered islands — San Cristobal — is extremely low (MacLeod et al., 2016). Underestimation is also common when several individuals are present in the image — generally the more individuals per image, the lower the agreement between CS and the experts (Hines et al., 2015; Jones et al., 2018). However, our statistical analyses showed that volunteers can also count accurately when several iguanas are present, except for in phase 1.

### Can citizen scientists accurately identify and count marine iguanas on a given aerial image?

In order to address this question, we must consider both the participation rate of the volunteers — to assess whether analysing a dataset within a reasonable timeframe is likely — and the accuracy of the volunteer-generated data.

In our study, we used the online citizen science platform ‘Zooniverse’, which was established in 2007 (Lintott et al., 2008). Astronomy-related projects initially dominated the platform, however, recent years have seen a shift towards projects with a focus on Nature and Biology (Cox et al., 2015; Zooniverse.org website). The platform has hosted over 400 projects and has more than 2.5 million registered volunteers (Lawson et al., 2022) who — along with non-registered users — are free to participate in any project lodged there. In our cumulative curve of participation, we experienced a daily classification rate of 500-1500, which increased in response to promotional work; in keeping with results from other projects (Lintott et al., 2008; Spiers et al., 2019). Crowdsourcing undoubtedly offers the opportunity to analyse large datasets with relatively little expert input, but without active promotion and engagement, it can take considerably longer to complete the analysis (Hennon et al., 2015). Due to time constraints within our project, promotional work was limited, therefore our project took longer to complete than other high profile Zooniverse projects (e.g. Snapshot Serengeti, which received 1 million classifications within 3 days of launch, or the Galaxy Zoo which received up to 70,000 classifications per hour). Each phase of our project took between 5 and 14 months to complete. This timeframe was influenced by the size of the phase dataset, with an overall average of almost 1900 images being fully analysed each month (Phase 1: 2031; Phase 2: 1819; Phase 3: 1740). However, for our purposes, this length of time was manageable, and there are clear ways to speed up the process via promotion if time is pressing.

After applying the best-performing aggregation method to the CS data, and omitting data from the pilot phase of the project — where data collection was suboptimal — we find that CS-counts were 90% and 92% accurate when compared to those of experts. This meets the criteria we defined as “accurate” in terms of counts, and thus we find the approach suitable for the task of counting marine iguanas. Although expert counts in phase 2 and 3 are not significantly higher than those from the CS data, a tendency for volunteers to underestimate is still evident. This indicates the need to calculate and apply a correction factor if CS inputs are to be used for the purpose of calculating population size. We are currently running phase 4 of the project and intend to use these data alongside those already collected — which in total will cover all types of terrain and colony density — for this purpose.

Since CS projects are completely reliant on volunteer inputs, it is important to consider the volunteer experience and factors that motivate participants. A study by Aceves-Bueno et al. (2017) — which analysed projects in which citizen scientists collected data — found that participants who receive economic recognition outperformed those who did not. Whilst this type of ‘reward’ may be helpful, our experience here indicates that recognition for work undertaken is not a prerequisite for involvement. Our finding — that even anonymous volunteers made important contributions — is in keeping with that of similar projects, where many participating volunteers remained anonymous (not registered; Jones et al., 2018; Shamir et al., 2014). It seems that volunteers in these types of projects tend to not expect external rewards, but may rather be motivated by the desire to contribute to science/conservation, engage with researchers, and get involved in scientific discussion (Mugar et al., 2014). Since the only training such volunteers obtain is that provided online by the project, and in discussions undertaken with the researchers, anyone with access to the internet, no matter their location or educational background, can access the data, making involvement in such projects easily accessible to a broad range of people. Another pattern noted in previous work, which was confirmed in our project, is that most participants contribute few classifications, while few volunteers contribute many (Anton et al., 2018; Hennon et al., 2015; Kiskin et al., 2020). In our case we found several volunteers that classified several thousand images, including one that looked at all images within the phase. This finding was confirmed by the volunteers’ replies to our survey, where most users responding estimated themselves to have contributed up to 100 classifications. This finding makes a strong case for the use of multiple independent classifications for each image; since this allows a dataset to be rapidly analysed even when the contribution of each user is small.

It is also important to consider factors that reduce volunteer motivation; in our case, the infrequency of marine iguanas in the images seems to have been important, as reported by the volunteers (see Supplementary Fig. S8). Interestingly, other researchers have found that such ‘blank’ images motivated the volunteers to keep looking for the target, and when blanks were removed, their participation time decreased (Bowyer et al., 2015); though worth noting is that this study did not explicitly test number of classifications made in relation to proportion of blank images. In our dataset, where 90% of our images were blank, volunteers contributing only a few classifications may not have seen any iguanas, and may have also spent a large amount of time attempting to distinguish these objects from a visually similar substrate. It would be interesting to investigate how filtering of blanks from our dataset would affect the outcomes, and we plan to investigate the utility of Machine Learning to automate the removal of such images.

### Consideration of online citizen science projects

Image-based datasets for wildlife monitoring are increasingly used, in great part due to technological advances that have made devices — such as camera traps and drones — more affordable and more suitable for such work. However, although these approaches can reduce survey time, effort required for image analysis can constitute a considerable burden for projects with limited resources. Online citizen science projects — which involve collaboration of the general public in order to analyse images remotely — are helping to resolve this issue, offering key advantages such as increased cost-effectiveness and a significant reduction in workload.

Despite the historical debate regarding the accuracy of citizen science-generated data (Aceves-Bueno et al., 2017); online CS is now recognized as an important approach for the undertaking of large-scale ecological research (Dickinson et al., 2010; McShea et al., 2016). Multiple studies have found that researchers can obtain accurate data from volunteers by properly aggregating multiple independent responses for one subject (e.g. an image or audio file; Anton et al., 2018; Egna et al., 2020; Swanson et al., 2016; Torney et al., 2019). Recent research has identified the number of independent classifications required to give accurate results, thereby streamlining the approach whilst still obtaining scientifically valid outcomes (Swanson et al., 2015).

One big draw of the CS approach is that by crowd-sourcing the collection and analysis of data, researchers can meet their aims in less time and/or expand their aims past what would be possible using more traditional approaches (Spiers et al., 2019; Willi et al., 2019). A review across 17 CS projects estimated that the CS approach allowed analysis to be completed on average within 2.4 years per project, as opposed to the 37 years estimated if the data were classified by experts (Cox et al., 2015). Consequently, in addition to meeting initial goals of projects — such as the identification and quantification of individuals — scientist using CS have been able to study further questions using their datasets (Beaudrot et al., 2020; Cosentino & Gibbs, 2022; Houskeeper et al., 2022; McCarthy et al., 2021; Nowak et al., 2020).

Large-scale monitoring in the Galápagos is logistically extremely challenging and is therefore rare (Páez-Rosas et al., 2021; Vargas et al., 2005). For most of the species, the majority of studies focus on just a few colonies (Ortiz-Catedral et al., 2023; Valle, 1995). For the marine iguana, surveying the whole range of the species is only realistic when new approaches — such as drone-based surveying — are applied, but analysis of the large datasets generated currently remains a significant obstacle to completion of this work. Here we confirm that aerial images have the potential to provide reliable data from volunteers with little training, indicating a reasonable approach to address the analysis bottleneck. Moreover, the images can be used to address numerous questions related to other taxa and the environment. Apart from the tasks related to the marine iguanas, we also asked the volunteers to classify other species and detect plastic objects; this is data we were able to easily collect alongside our tasks, which will be made available to other researchers as a contribution to their work. We find that the combination of an image-based approach in surveying remote areas, together with crowdsourced analysis via online CS, can be a valuable approach in the collection and analysis of large datasets.

In parallel, researchers may also use CS to engage and educate the public on themes related to science and conservation (Kosmala et al., 2016). Studies have shown that citizen scientists who participate in projects tend to develop positive environmental attitudes as a result (Bonney et al., 2016). Therefore, involving the public may not only benefit the project in question, but may also help the scientific community, aid in public engagement and education, and could increase interest in topics related to biodiversity and the environment.

## Conclusion and further work

Our results validate the use of the citizen science approach to accurately identify and count marine iguanas from aerial images, however, there is a tendency to underestimate the number of iguanas.

Our next step is to continue our analysis which includes images from the most populated colonies, in order to identify a correction factor that will allow CS inputs to be used for accurate population-size estimates of marine iguanas. This is possible because the counting of iguanas from aerial imagery by experts has already been validated against traditional ground-based approaches (Varela-Jaramillo et al., 2023). This work is an important contribution to our overall goal of addressing the population-size data-gap that currently hampers effective conservation in this species (MacLeod et al., 2020).

In future work, we expect to use our images to also analyse reproduction dynamics, and potentially habitat characteristics, using a CS approach. We are interested to see whether volunteers can reliably identify certain aspects of the colonies, such as the presence of leks and males with breeding coloration; these data will be useful to address the dearth of information regarding marine iguana breeding activity across the archipelago.

Our next major goal is to use machine learning (ML) to analyze drone imagery. As with several other projects (Bird et al., 2018; Jones et al., 2020; Torney et al., 2019; Willi et al., 2019), CS-input is being used to train Artificial Intelligence for pattern recognition and minimize training time for the computers. We also expect to use ML to filter images, enabling us to remove blank images from our CS datasets so that we can focus this human effort on the images where it is most needed. We envisage that this will improve volunteer participation and decrease the running time of the online project. By combining CS with ML, we ultimately aim to create a semi-automated pipeline capable of finding and counting marine iguanas and other biologically relevant objects in drone images, thereby greatly reducing the amount of effort needed to undertake such work.

## Acknowledgements

We thank all our Zooniverse volunteers, especially those that were particularly loyal to the project and contributed significant amounts of time and effort. A special mention must be made to our most loyal and qualified volunteer, Pamela Van Schouwen (@Pamelavans at Zooniverse) who also helped us as moderator over two phases of the project. We thank the Galápagos National Park and the Ministerio de Ambiente, Agua y Transición Ecológica de Ecuador for granting permits (No. PC-57-20, PCD-71-21 and PC-90-22). We are thankful to the Galápagos Science Center staff for helping with field logistics. We are particularly grateful to Andrés Mármol (3Diversity) for advice on data analysis, to Efrain Reveló, Manuel Yépez (Sharksky Galápagos Travel & Conservation) and Lenin Cruz for providing safe transportation. We thank Gustavo Pazmiño, Kathleen Preissler, Denisse Dalgo & Sophie Timmerman for helping with data collection. Fieldwork was supported by the University of Leipzig via start-up funding for the research group of SS at the faculty of Natural Science. We are grateful for the additional funds granted to AM by the International Iguana Foundation, the Swiss Friends of the Galápagos, and the Galapagos Conservation Trust. Research by JMG and GRT is supported by Universidad San Francisco de Quito and the Galapagos Science Center. AV is supported by a stipend from the German Academic Exchange Service.

## Notes

https://doi.org/10.6084/m9.figshare.25196306

### Competing Interest Statement

The authors have declared no competing interest.

## Literature

Aceves-Bueno, E., Adeleye, A. S., Feraud, M., Huang, Y., Tao, M., Yang, Y., & Anderson, S. E. (2017). The Accuracy of Citizen Science Data: A Quantitative Review. The Bulletin of the Ecological Society of America, 98(4), 278–290. 10.1002/bes2.1336

Alava, J. J., McMullen, K., Jones, J., Barragán-Paladines, M. J., Hobbs, C., Tirapé, A., Calle, P., Alarcón, D., Muñoz-Pérez, J. P., Muñoz-Abril, L., Townsend, K. A., Denkinger, J., Uyaguari, M., Domínguez, G. A., Espinoza, E., Reyes, H., Piedrahita, P., Fair, P., Galloway, T., … Schofield, J. (2023). Multiple anthropogenic stressors in the Galápagos Islands’ complex social–ecological system: Interactions of marine pollution, fishing pressure, and climate change with management recommendations. Integrated Environmental Assessment and Management, 19(4), 870–895. 10.1002/ieam.4661

Anton, V., Hartley, S., Geldenhuis, A., & Wittmer, H. U. (2018). Monitoring the mammalian fauna of urban areas using remote cameras and citizen science. Journal of Urban Ecology, 4(1). 10.1093/jue/juy002

Beaudrot, L., Palmer, M. S., Anderson, T. M., & Packer, C. (2020). Mixed-species groups of Serengeti grazers: A test of the stress gradient hypothesis. Ecology, 101(11), e03163. 10.1002/ecy.3163

Berger-Wolf, T. Y., Rubenstein, D. I., Stewart, C. V., Holmberg, J. A., Parham, J., Menon, S., Crall, J., Van Oast, J., Kiciman, E., & Joppa, L. (2017). *Wildbook: Crowdsourcing, computer vision, and data science for conservation* (arXiv:1710.08880). arXiv. 10.48550/arXiv.1710.08880

Bird, R., Daniel, M., Dickinson, H., Feng, Q., Fortson, L., Furniss, A., Jarvis, J., Mukherjee, R., Ong, R., Sadeh, I., & Williams, D. (2018). Muon Hunter: A Zooniverse project. Journal of Physics: Conference Series, 1342. 10.1088/1742-6596/1342/1/012103

Bland, L. M., Bielby, J., Kearney, S., Orme, C. D. L., Watson, J. E. M., & Collen, B. (2017). Toward reassessing data-deficient species. Conservation Biology, 31(3), 531–539.

Bonney, R., Phillips, T. B., Ballard, H. L., & Enck, J. W. (2016). Can citizen science enhance public understanding of science? Public Understanding of Science, 25(1), 2–16. 10.1177/0963662515607406

Bowyer, A., Maidel, V., Lintott, C,, Swanson, A., & Miller, G.. (2015). This Image Intentionally Left Blank: Mundane Images Increase Citizen Science Participation. 10.13140/RG.2.2.35844.53121

Cardoso, A., Malhi, Y., Oliveras, I., Lehmann, D., Ndong, J., Dimoto, E., Bush, E., Jeffery, K., Labriere, N., Lewis, S., White, L., Bond, W., & Abernethy, K. (2020). The Role of Forest Elephants in Shaping Tropical Forest–Savanna Coexistence. Ecosystems, 23. 10.1007/s10021-019-00424-3

Cosentino, B. J., & Gibbs, J. P. (2022). Parallel evolution of urban–rural clines in melanism in a widespread mammal. Scientific Reports, 12(1), Article 1. 10.1038/s41598-022-05746-2

Cox, J., Oh, E. Y., Simmons, B., Lintott, C., Masters, K., Greenhill, A., Graham, G., & Holmes, K. (2015). Defining and Measuring Success in Online Citizen Science: A Case Study of Zooniverse Projects. Computing in Science & Engineering, 17(4), 28–41. 10.1109/MCSE.2015.65

Dickinson, J. L., Zuckerberg, B., & Bonter, D. N. (2010). Citizen Science as an Ecological Research Tool: Challenges and Benefits. *Annual Review of Ecology*, Evolution, and Systematics, 41(1), 149–172. 10.1146/annurev-ecolsys-102209-144636

Egna, N., O’Connor, D., Stacy-Dawes, J., Tobler, M. W., Pilfold, N., Neilson, K., Simmons, B., Davis, E. O., Bowler, M., Fennessy, J., Glikman, J. A., Larpei, L., Lekalgitele, J., Lekupanai, R., Lekushan, J., Lemingani, L., Lemirgishan, J., Lenaipa, D., Lenyakopiro, J., … Owen, M. (2020). Camera settings and biome influence the accuracy of citizen science approaches to camera trap image classification. Ecology and Evolution, 10(21), 11954–11965. 10.1002/ece3.6722

Ezat, M. A., Fritsch, C. J., & Downs, C. T. (2018). Use of an unmanned aerial vehicle (drone) to survey Nile crocodile populations: A case study at Lake Nyamithi, Ndumo game reserve, South Africa. Biological Conservation, 223, 76–81. 10.1016/j.biocon.2018.04.032

Hennon, C. C., Knapp, K. R., Schreck, C. J., Stevens, S. E., Kossin, J. P., Thorne, P. W., Hennon, P. A., Kruk, M. C., Rennie, J., Gadéa, J.-M., Striegl, M., & Carley, I. (2015). Cyclone Center: Can Citizen Scientists Improve Tropical Cyclone Intensity Records? Bulletin of the American Meteorological Society, 96(4), 591–607. 10.1175/BAMS-D-13-00152.1

Hines, G., Swanson, A., Kosmala, M., & Lintott, C. (2015). Aggregating User Input in Ecology Citizen Science Projects. Proceedings of the AAAI Conference on Artificial Intelligence, 29(2), Article 2. 10.1609/aaai.v29i2.19057

Houskeeper, H. F., Rosenthal, I. S., Cavanaugh, K. C., Pawlak, C., Trouille, L., Byrnes, J. E. K., Bell, T. W., & Cavanaugh, K. C. (2022). Automated satellite remote sensing of giant kelp at the Falkland Islands (Islas Malvinas). PLOS ONE, 17(1), e0257933. 10.1371/journal.pone.0257933

IUCN. (2024). The IUCN Red List of Threatened Species. IUCN Red List of Threatened Species. https://www.iucnredlist.org/en

Jones, F. M., Allen, C., Arteta, C., Arthur, J., Black, C., Emmerson, L. M., Freeman, R., Hines, G., Lintott, C. J., Macháčková, Z., Miller, G., Simpson, R., Southwell, C., Torsey, H. R., Zisserman, A., & Hart, T. (2018). Time-lapse imagery and volunteer classifications from the Zooniverse Penguin Watch project. Scientific Data, 5(1), Article 1. 10.1038/sdata.2018.124

Jones, F. M., Arteta, C., Zisserman, A., Lempitsky, V., Lintott, C. J., & Hart, T. (2020). Processing citizen science-and machine-annotated time-lapse imagery for biologically meaningful metrics. Scientific Data, 7(1), Article 1. 10.1038/s41597-020-0442-6

Kiskin, I., Cobb, A. D., Wang, L., & Roberts, S. (2020). Humbug Zooniverse: A Crowd-Sourced Acoustic Mosquito Dataset. 2020*-May*, 916-920. 10.1109/ICASSP40776.2020.9053141

Kosmala, M., Wiggins, A., Swanson, A., & Simmons, B. (2016). Assessing data quality in citizen science. Frontiers in Ecology and the Environment, 14, 551–560. 10.1002/fee.1436

Lawson, K. N., Tracy, B. M., Sharova, M., Muirhead, J. R., & Cawood, A. (2022). Engaging Online Citizen Scientists and the Consensus Method to Monitor the Marine Biofouling Community. Frontiers in Marine Science, 9. 10.3389/fmars.2022.862430

Lee, W. Y., Park, M., & Hyun, C.-U. (2019). Detection of two Arctic birds in Greenland and an endangered bird in Korea using RGB and thermal cameras with an unmanned aerial vehicle (UAV). PLOS ONE, 14(9), e0222088. 10.1371/journal.pone.0222088

Lenth, R. V., Bolker, B., Buerkner, P., Giné-Vázquez, I., Herve, M., Jung, M., Love, J., Miguez, F., Riebl, H., & Singmann, H. (2023). *emmeans: Estimated Marginal Means, aka Least-Squares Means* (1.8.9). https://cran.r-project.org/web/packages/emmeans/index.html

Lintott, C. J., Schawinski, K., Slosar, A., Land, K., Bamford, S., Thomas, D., Raddick, M. J., Nichol, R. C., Szalay, A., Andreescu, D., Murray, P., & Berg, J. van den. (2008). Galaxy Zoo: Morphologies derived from visual inspection of galaxies from the Sloan Digital Sky Survey. Monthly Notices of the Royal Astronomical Society, 389(3), 1179–1189. 10.1111/j.1365-2966.2008.13689.x

MacLeod, A., Nelson, K. N., & Grant, T. D. (2020). *Amblyrhynchus cristatus*. IUCN Red List of Threatened Species. https://www.iucnredlist.org/species/1086/177552193

MacLeod, A., & Steinfartz, S. (2016). The conservation status of the Galápagos marine iguanas, Amblyrhynchus cristatus: A molecular perspective. Amphibia Reptilia, 37(1), 91–109. 10.1163/15685381-00003035

MacLeod, A., Unsworth, L., Trillmich, F., & Steinfartz, S. (2016). Mark-resight estimates confirm a critically small population size in threatened marine iguanas (Amblyrhynchus cristatus) on San Cristóbal Island, Galápagos. Salamandra, 52(1), 58–62.

Marshall, N. J., Kleine, D. A., & Dean, A. J. (2012). CoralWatch: Education, monitoring, and sustainability through citizen science. Frontiers in Ecology and the Environment, 10(6), 332–334. 10.1890/110266

McCarthy, M. S., Stephens, C., Dieguez, P., Samuni, L., Després-Einspenner, M.-L., Harder, B., Landsmann, A., Lynn, L. K., Maldonado, N., Ročkaiová, Z., Widness, J., Wittig, R. M., Boesch, C., Kühl, H. S., & Arandjelovic, M. (2021). Chimpanzee identification and social network construction through an online citizen science platform. Ecology and Evolution, 11(4), 1598–1608. 10.1002/ece3.7128

McShea, W. J., Forrester, T., Costello, R., He, Z., & Kays, R. (2016). Volunteer-run cameras as distributed sensors for macrosystem mammal research. Landscape Ecology, 31(1), 55–66. 10.1007/s10980-015-0262-9

Miralles, A., Macleod, A., Rodríguez, A., Ibáñez, A., Jiménez-Uzcategui, G., Quezada, G., Vences, M., & Steinfartz, S. (2017). Shedding light on the Imps of Darkness: An integrative taxonomic revision of the Galápagos marine iguanas (genus Amblyrhynchus). Zoological Journal of the Linnean Society, 181(3), 678–710. 10.1093/zoolinnean/zlx007

Monks, J., Wills, H., & Knox, C. (2022). Testing Drones as a Tool for Surveying Lizards. Drones, 6, 199. 10.3390/drones6080199

Mugar, G., Østerlund, C., Hassman, K., Crowston, K., & Jackson, C. (2014). Planet hunters and seafloor explorers: Legitimate peripheral participation through practice proxies in online citizen science. En *Proceedings of the ACM Conference on Computer Supported Cooperative Work*, CSCW (p. 119). 10.1145/2531602.2531721

Nowak, K., Berger, J., Panikowski, A., Reid, D. G., Jacob, A. L., Newman, G., Young, N. E., Beckmann, J. P., & Richards, S. A. (2020). Using community photography to investigate phenology: A case study of coat molt in the mountain goat (Oreamnos americanus) with missing data. Ecology and Evolution, 10(23), 13488–13499. 10.1002/ece3.6954

Ortiz-Catedral, L., Kumar, K., Llerena, A. J., Jie, C. H. Z., Sollis, H., Ramirez, J., Gavilanes, M., Chimborazo, W., Guerrero, B., Sevilla, C., & Rueda, D. (2023). <p>Life on the volcano: Population size of the Galápagos Land Iguana <em>Conolophus subcristatus</em> (Gray, 1831) on Fernandina Island, Galápagos, Ecuador</p>. Herpetology Notes, 16, 41–47.

Páez-Rosas, D., Torres, J., Espinoza, E., Marchetti, A., Seim, H., & Riofrío-Lazo, M. (2021). Declines and recovery in endangered Galapagos pinnipeds during the El Niño event. Scientific Reports, 11(1), Article 1. 10.1038/s41598-021-88350-0

Pedregosa, F., Varoquaux, G., Gramfort, A., Michel, V., Thirion, B., Grisel, O., & others. (2011). Scikit-learn: Machine learning in Python. Journal of Machine Learning Research, 2825–2830.

Ratcliffe, N., Guihen, D., Robst, J., Crofts, S., Stanworth, A., & Enderlein, P. (2015). A protocol for the aerial survey of penguin colonies using UAVs. Journal of Unmanned Vehicle Systems, 3(3), 95–101. 10.1139/juvs-2015-0006

RStudio Team. (2023). *RStudio: Integrated Development for R. RStudio* [Software]. http://www.rstudio.com/

Shamir, L., Yerby, C., Simpson, R., von Benda-Beckmann, A. M., Tyack, P., Samarra, F., Miller, P., & Wallin, J. (2014). Classification of large acoustic datasets using machine learning and crowdsourcing: Application to whale calls. The Journal of the Acoustical Society of America, 135(2), 953–962. 10.1121/1.4861348

Simpson, R., Page, K. R., & De Roure, D. (2014). Zooniverse: Observing the world’s largest citizen science platform. 1049–1054. 10.1145/2567948.2579215

Spiers, H., Swanson, A., Fortson, L., Simmons, B. D., Trouille, L., Blickhan, S., & Lintott, C. (2019). Everyone counts? Design considerations in online citizen science. Journal of Science Communication, 18(1). 10.22323/2.18010204

Sullivan, B. L., Aycrigg, J. L., Barry, J. H., Bonney, R. E., Bruns, N., Cooper, C. B., Damoulas, T., Dhondt, A. A., Dietterich, T., Farnsworth, A., Fink, D., Fitzpatrick, J. W., Fredericks, T., Gerbracht, J., Gomes, C., Hochachka, W. M., Iliff, M. J., Lagoze, C., La Sorte, F. A., … Kelling, S. (2014). The eBird enterprise: An integrated approach to development and application of citizen science. Biological Conservation, 169, 31–40. 10.1016/j.biocon.2013.11.003

Swanson, A., Kosmala, M., Lintott, C., & Packer, C. (2016). A generalized approach for producing, quantifying, and validating citizen science data from wildlife images. Conservation Biology, 30(3), 520–531. 10.1111/cobi.12695

Swanson, A., Kosmala, M., Lintott, C., Simpson, R., Smith, A., & Packer, C. (2015). Snapshot Serengeti, high-frequency annotated camera trap images of 40 mammalian species in an African savanna. Scientific Data, 2. 10.1038/sdata.2015.26

The Pandas Development Team. (2020). Pandas version 2.1.2 [Software]. Zenodo. 10.5281/zenodo.3509134

Torney, C. J., Lloyd-Jones, D. J., Chevallier, M., Moyer, D. C., Maliti, H. T., Mwita, M., Kohi, E. M., & Hopcraft, G. C. (2019). A comparison of deep learning and citizen science techniques for counting wildlife in aerial survey images. Methods in Ecology and Evolution, 10(6), 779–787. 10.1111/2041-210X.13165

Valle, C. A. (1995). Effective Population Size and Demography of the Rare Flightless Galapagos Cormorant. Ecological Applications, 5(3), 601–617. 10.2307/1941970

Van Rossum, G., & Drake, F. L. (2009). Python 3 Reference Manual. Scotts Valley, *CA*: *CreateSpace*.

Varela-Jaramillo, A., Rivas-Torres, G., Guayasamin, J. M., Steinfartz, S., & MacLeod, A. (2023). A pilot study to estimate the population size of endangered Galápagos marine iguanas using drones. Frontiers in Zoology, 20(1), 4. 10.1186/s12983-022-00478-5

Vargas, H., Lougheed, C., & Snell, H. (2005). Population size and trends of the Galápagos Penguin Spheniscus mendiculus. Ibis, 147(2), 367–374. 10.1111/j.1474-919x.2005.00412.x

Willi, M., Pitman, R. T., Cardoso, A. W., Locke, C., Swanson, A., Boyer, A., Veldthuis, M., & Fortson, L. (2019). Identifying animal species in camera trap images using deep learning and citizen science. Methods in Ecology and Evolution, 10(1), 80–91. 10.1111/2041-210X.13099

Wood, S. A., Robinson, P. W., Costa, D. P., & Beltran, R. S. (2021). Accuracy and precision of citizen scientist animal counts from drone imagery. PLOS ONE, 16(2), e0244040. 10.1371/journal.pone.0244040

