## Supplementary Materials for "Iguanas from above: Citizen scientists provide reliable counts of endangered Galápagos marine iguanas from drone imagery"

**SUPPLEMENTARY INFORMATION**

### 1. Supplementary Figures

**Figure S1.** Example image representing the number of iguanas counted by the experts compared to all the volunteers' marks given to the same image. The spatial marks were used by the HDBSCAN clustering method to estimate a number of spatial clusters, which represent the aggregated number of iguanas counted by the volunteers. Here we set a minimum sample of 5 marks to form a cluster.

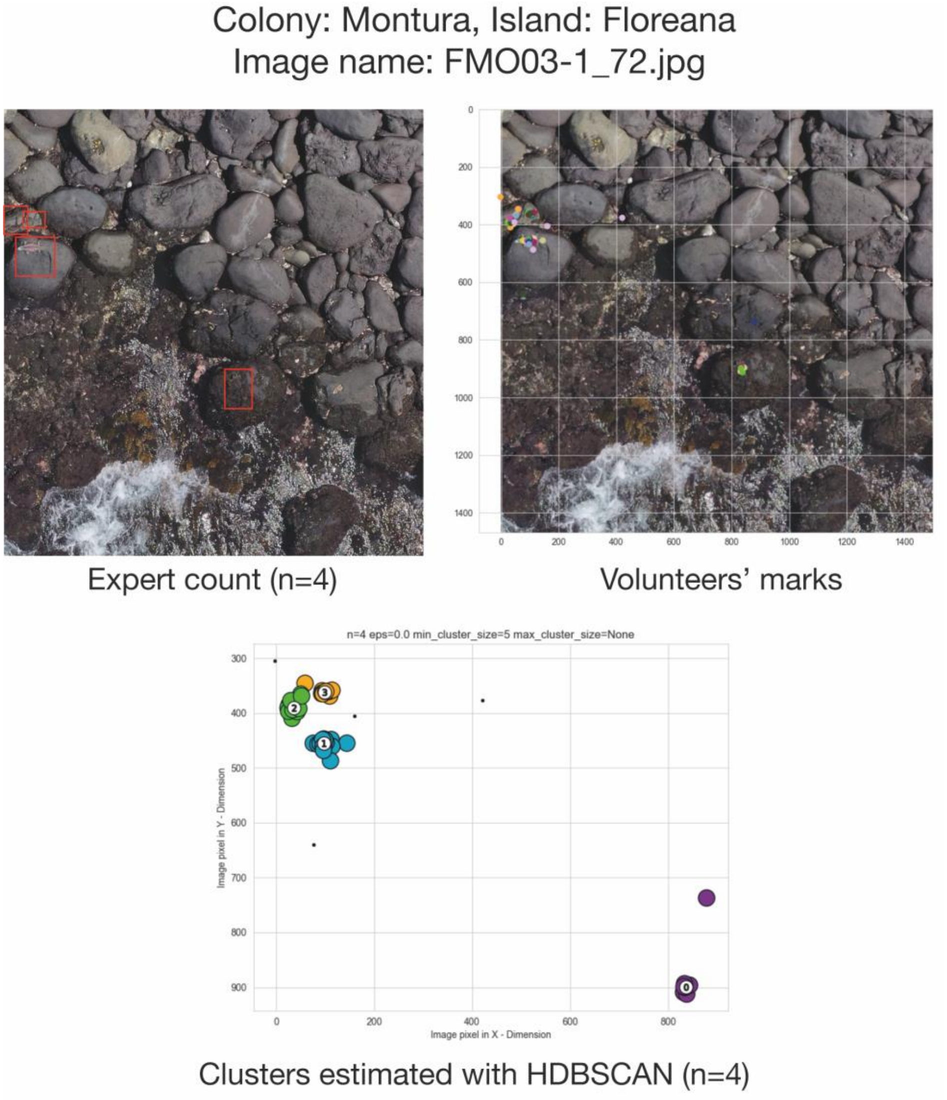

**Figure S2.** Plot of marine iguanas counted in the GS dataset. Comparison of expert counts vs CS counts aggregated with the HDBSCAN clustering method.

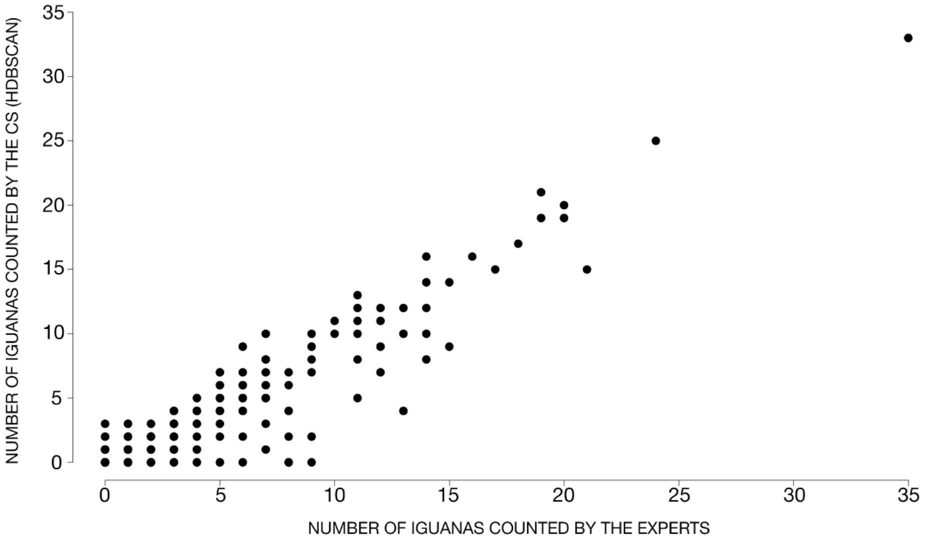

**Figure S3.** Distribution graph of the number of classifications done by all the citizen scientists who participated along our three phases (A) and the estimated number of classifications done by 110 volunteers that replied to our survey (B).

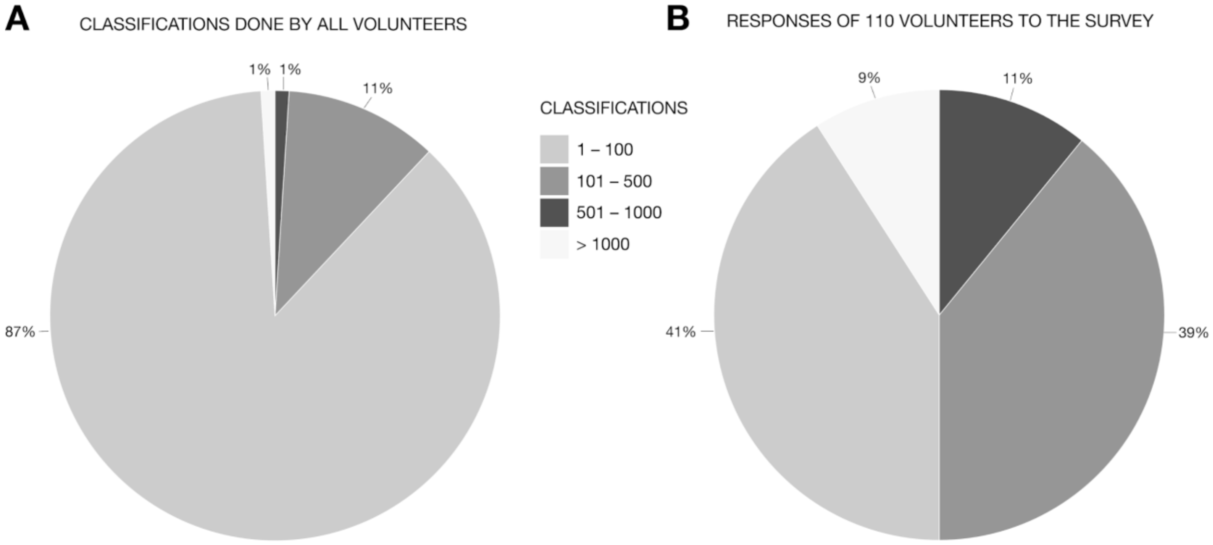

**Figure S4.** Results regarding CS accuracy in the GS dataset (on images with iguanas present and absent) when using the majority vote rule to assess if an answer was correct or incorrect after comparing it against the expert answer.

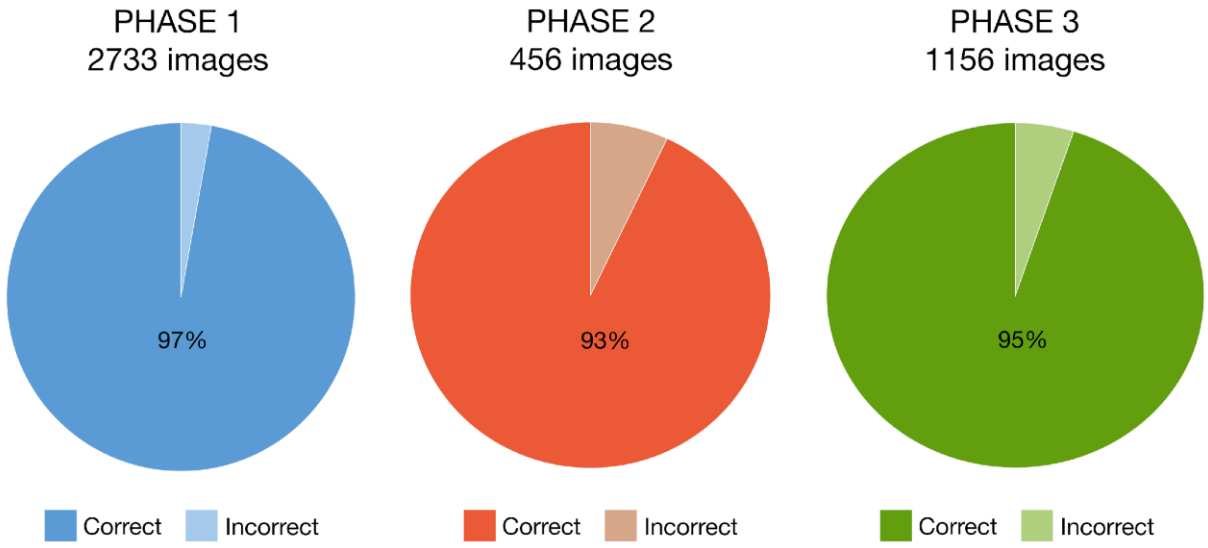

**Figure S5.** Plots of the generalized linear models presented independently by phase when the factor quality of the image is added to the analysis, assessing differences amongst methods used to count marine iguanas from the GS dataset.

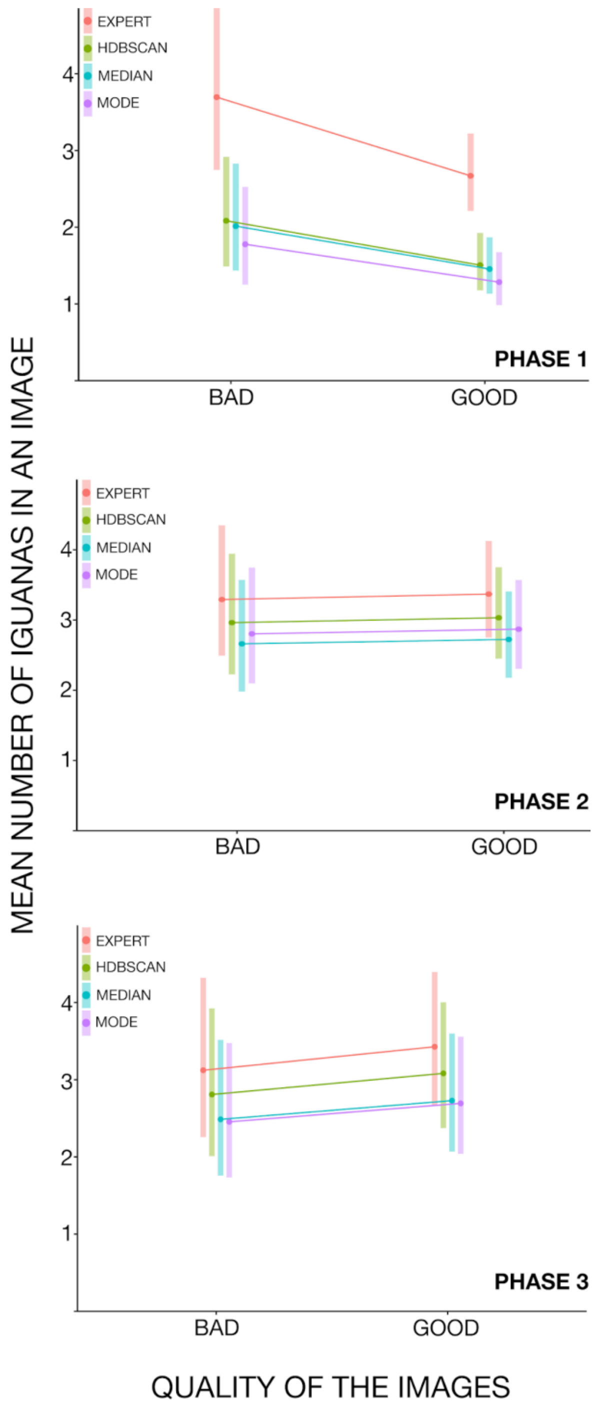

**Figure S6.** Plots of the generalized linear models presented independently by phase when the factor 'number of iguanas present in the image' is added to the analysis, assessing differences amongst methods used to count marine iguanas from the GS dataset.

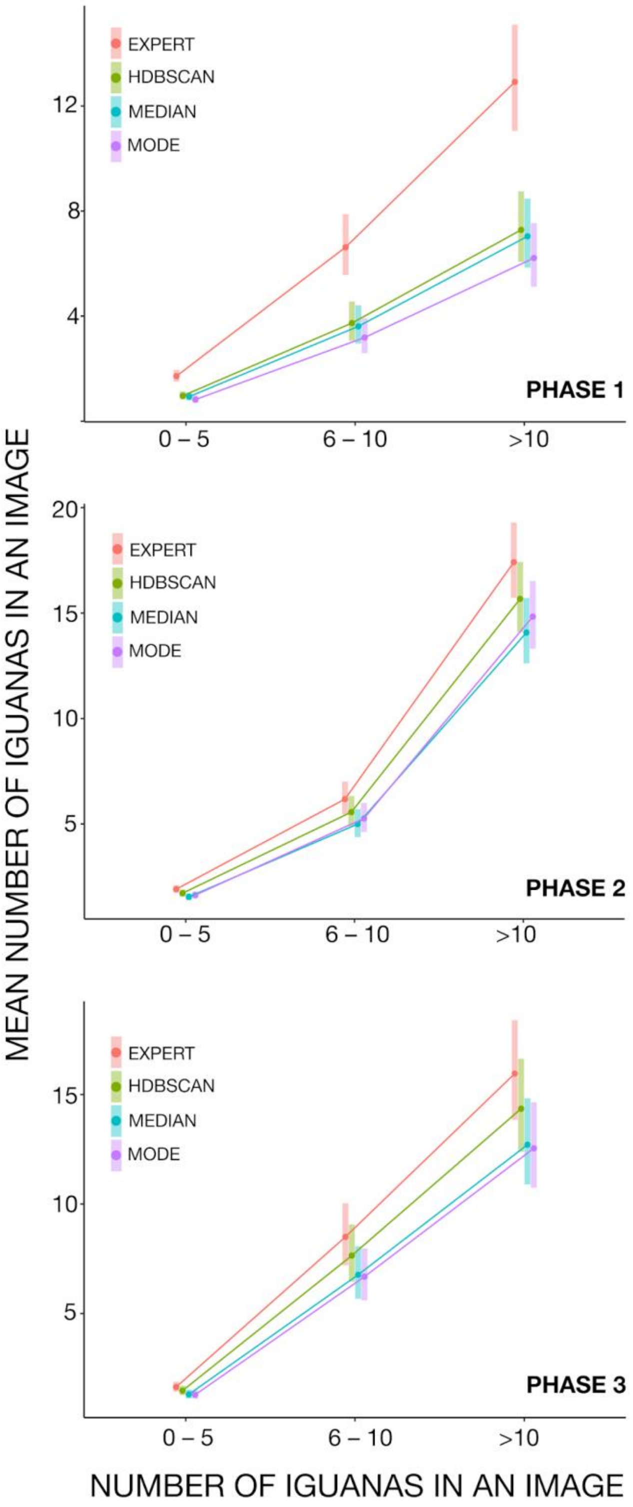

**Figure S7.** Survey results of 110 volunteers regarding self-assessed difficulties to count marine iguanas in the images, in relation to quality of the image and number of iguanas present in the image.

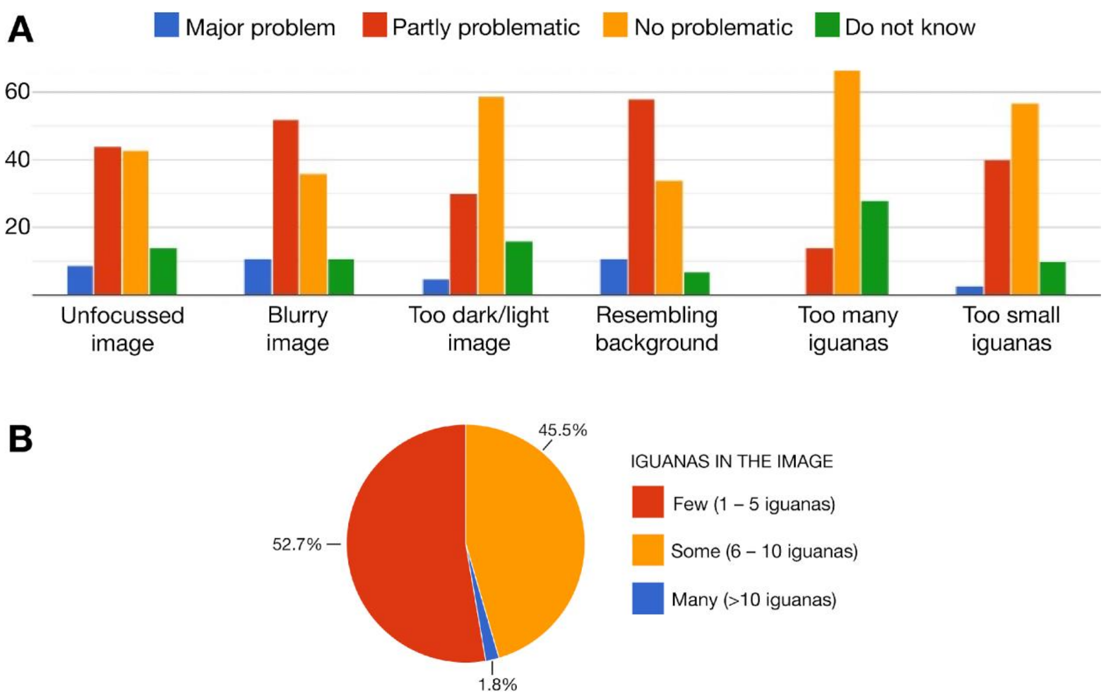

**Figure S8.** Survey results of 110 volunteers regarding factors affecting their motivation to contribute further to the CS project.

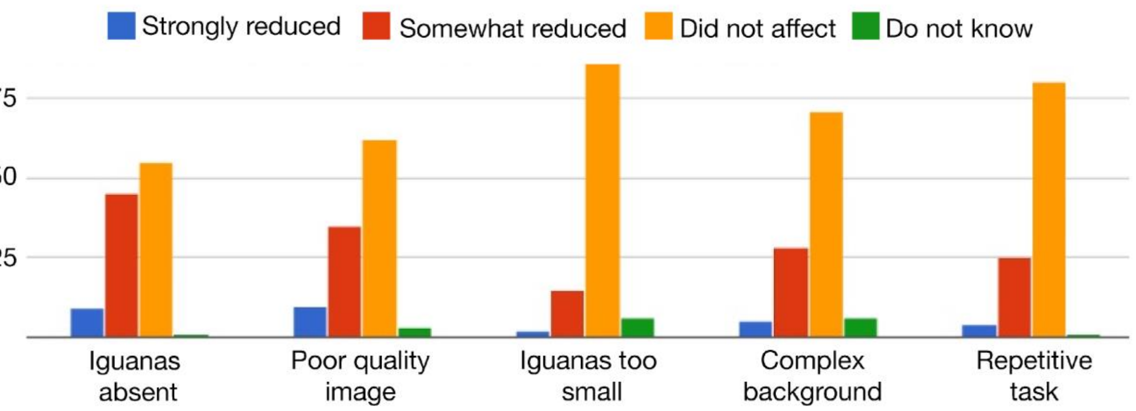

### 2. Supplementary Tables

**Table S1.** Number of images where CS counted equal to the experts, CS counted less than the experts and CS counted more than the experts. Values in bold represent best results.

| Phase (GS images) | Metric | CS = experts | | CS < experts (underestimation) | | CS > experts (overestimation) | |
|---|---|---|---|---|---|---|---|
| | | # images | % | # images | % | # images | % |
| All (4345) | Median | 4095 | 94.2 | 204 | 4.7 | 46 | 1.0 |
| | Mode | 4105 | 94.5 | 191 | 4.4 | 49 | **1.1** |
| | **HDBSCAN** | 4111 | **94.6** | 178 | **4.1** | 56 | 1.3 |
| | DBSCAN | 4003 | 92.1 | 256 | 5.9 | 86 | 2.0 |
| 1st (2733) | Median | 2646 | 96.8 | 76 | 2.8 | 11 | 0.4 |
| | **Mode** | 2647 | **96.9** | 77 | **2.8** | 9 | **0.3** |
| | HDBSCAN | 2644 | 96.7 | 76 | **2.8** | 13 | 0.5 |
| | DBSCAN | 2602 | 95.2 | 106 | 3.9 | 25 | 0.9 |
| 2nd (456) | Median | 360 | 78.9 | 73 | 16 | 23 | **5** |
| | Mode | 368 | 80.7 | 63 | 13.8 | 25 | 5.5 |
| | **HDBSCAN** | 374 | **82.0** | 58 | **12.7** | 24 | 5.3 |
| | DBSCAN | 337 | 73.9 | 82 | 18 | 37 | 8.1 |
| 3rd (1156) | Median | 1085 | 93.9 | 55 | 4.8 | 11 | **1.0** |
| | Mode | 1086 | 93.9 | 51 | 4.4 | 14 | 1.2 |
| | **HDBSCAN** | 1089 | **94.2** | 44 | **3.8** | 18 | 1.6 |
| | DBSCAN | 1060 | 91.7 | 68 | 5.9 | 23 | 2.0 |

**Table S2**. Results of the logistical regression used to find an R-square value which determines the highest fit (closest) between volunteers counts and expert counts, for all methods used to obtain volunteers counts.

| Method | Model | Nagelkerdes R2 | RMSE | Sigma | Score log | Performance score |
|---|---|---|---|---|---|---|
| HDBSCAN | glm | **0.912** | 3.403 | 1.215 | -1.954 | 96.9% |
| Median | glm | 0.885 | 3.691 | 1.339 | -2.055 | 49.86% |
| Mode | glm | 0.856 | 3.535 | 1.458 | -2.147 | 10.85% |

### 3. Supplementary Methods

#### 3.1. DBSCAN and HDBSCAN analyses.

First, marks were removed which depicted “partial iguanas” (i.e. individuals dissected as an artifact when images were sliced in image preparation). Next, images marked by fewer than 4 users were removed. Finally, we ran DBSCAN and HDBSCAN within the GS dataset where images were identified by experts to contain iguanas, using the minimum volunteer threshold, as follows:

1. DBSCAN relies on a minimum of two hyperparameters, eps for Epsilon, which is a threshold radius around any point. A point within the radius is considered a neighbour. Minimum Points refers to the points required to form a dense region.

   A grid search approach was chosen to find the best fitting parameter set, using Silhouette scoring as the metric for optimal fit. The search space was eps = [0.01, 0.05, 0.1, 0.2, 0.3, 0.4, 0.5], min_samples = [3, 5, 8, 10].

   Choosing eps and min_points effects found clusters. The optimal cluster count is found by maximizing the Silhouette score from each parameter. Two clusters are a minimum necessary to compare Silhouettes. This led to random results when sorting, when no two clusters could be found, resulting in worse outcomes.

2. HDBSCAN does not need an eps value to be set before the run. Default values were used, as min_cluster_sizes = 5 and cluster_selection_epsilon = 0.0.

Reference: Rousseeuw, P. J. 1987. Silhouettes: A graphical aid to the interpretation and validation cluster analysis. Journal of Computational and Applied Mathematics 20: 53–65. https://www.sciencedirect.com/science/article/pii/0377042787901257

**3.1.1.** Results of total counts per phase obtained with DBSCAN and HDBSCAN clustering methods with the root mean squared error (rmse) calculated when compared to the expert count.

| | **Phase 1 counts** | **Phase 1 rmse** | **Phase 2 counts** | **Phase 2 rmse** | **Phase 3 counts** | **Phase 3 rmse** |
|---|---|---|---|---|---|---|
| Expert count | 422 | - | 600 | - | 388 | - |
| DBSCAN count | 218 | 0.6441 | 482 | 1.2944 | 310 | 0.6322 |
| HDBSCAN count | 244 | 0.5694 | 541 | 0.6993 | 357 | 0.3891 |

**3.1.2.** Example from the colony: El Miedo, in Santa Fe Island, phase 1. Image showing all volunteers' marks and a table comparing the results of the different methods used (median, mode, DBSCAN and HDBSCAN) to aggregate data from volunteer classifications regarding the number of marine iguanas present in this image – these results are an indicator for why HDBSCAN (with minimum threshold of

5 volunteers marking iguanas(a)) generate better results, as the median selects the value in the middle of the sample while the mode choses the most repeated value.

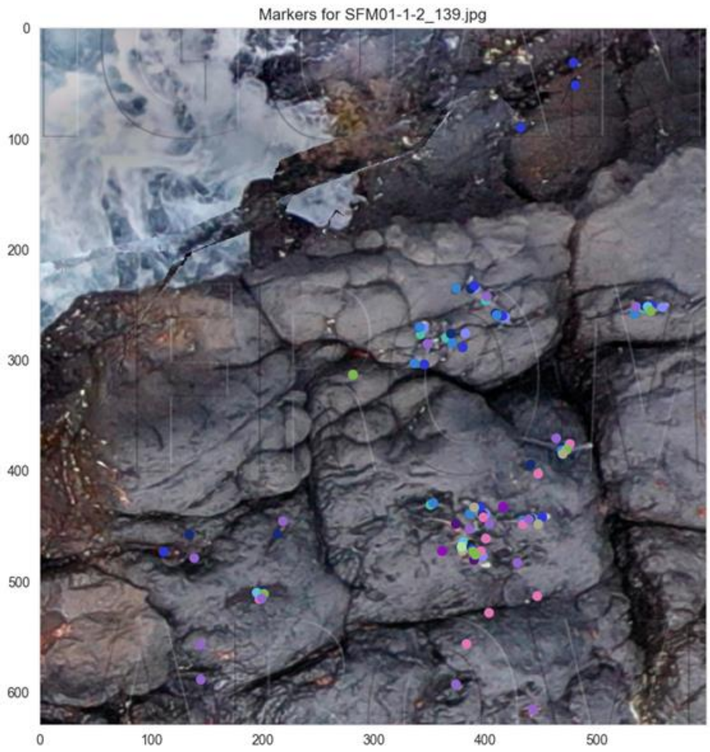


| Image name | CS counts in ascendent order | Median | Mode | DBSCAN | HDBSCAN | Expert count |
|---|---|---|---|---|---|---|
| SFM01-1-2_139.jpg | 1,1,1,2,2,4,4,5,9,9,10,11,12,14,15 | 5 | 1 | 7 | 8 | 11 |

#### 3.1.3. Code used to implement the DBSCAN and HDBSCAN clustering analyses into our data.

The Code is hosted on GitHub: https://github.com/cwinkelmann/iguanas-from-above-zooniverse

**Zooniverse data clustering for Iguanas from Above Project**

**3.2. Survey questions presented to our volunteers regarding their experiences within our project.**

**3.2.1. How many classifications** do you estimate you have made on this project?

- 1 – 100
- 101 – 500
- 501 – 1000
- > 1000

**3.2.2.** Please rate the following image features in terms of the **level of difficulty** they caused in finding the iguanas.

- Photo unfocused

- Image blurry/smeared
- Photo too dark/light
- Background too similar to iguanas
- Too many iguanas
- Iguanas too small

**3.2.3.** How much did the following aspects of the project **reduce your motivation** for classifying images?

- No iguanas in the images
- Poor quality image
- Iguanas too small
- Background too complex/similar to the iguanas
- Task repetitive

**3.2.4.** The iguanas were most **easily counted** in images with:

- Few iguanas (up to 5)
- Some iguanas (6–10)
- Many iguanas (> 11)
